# Milk losses and dynamics during perturbations in dairy cows differ with parity and lactation stage

**DOI:** 10.1101/2020.07.01.182568

**Authors:** I. Adriaens, I. van den Brulle, L. D’Anvers, J.M.E. Statham, K. Geerinckx, S. De Vliegher, S. Piepers, B. Aernouts

## Abstract

Milk yield dynamics during perturbations reflect how cows respond to challenges. This study investigated the characteristics of 62,406 perturbations from 16,604 lactation curves of dairy cows milked with an automated milking system at 50 Belgian, Dutch and English farms. The unperturbed lactation curve representing the theoretical milk yield dynamics was estimated with an iterative procedure fitting a Wood model on the daily milk yield data not part of a perturbation. Each perturbation was characterized and split in a development and a recovery phase. Based hereon, we calculated both the characteristics of the perturbation as a whole, and the duration, slopes and milk losses in the phases separately. A two-way analysis of variance followed by a pairwise comparison of group means was carried out to detect differences between these characteristics in different lactation stages (early, mid-early, mid-late and late) and parities (first, second and third or higher). On average, 3.8 ± 1.9 (mean ± standard deviation) perturbations were detected per lactation in the first 305 days after calving, corresponding to an estimated 92.1 ± 135.8 kg of milk loss. Only 1% of the lactations had no perturbations. The average development and recovery rates were respectively −2.3 and 1.5 kg per day, and these phases lasted on average 10.1 and 11.6 days. Perturbation characteristics were significantly different across parity and lactation stage groups, and early and mid-early perturbations in higher parities were found to be more severe, with faster development rates, slower recovery rates and higher milk losses. The method to characterize perturbations can be used for precision phenotyping purposes looking into the response of cows to challenges, or for monitoring applications, for example to evaluate the development and recovery of diseases and how these are affected by preventive actions or treatments.

## INTRODUCTION

Milk yield and production performance of dairy cows are the result of both genetic and environmental factors. In the past decades, substantial progress has been made in terms of genetic predisposition for milk yield through dedicated selection, and via the increased use of advanced breeding tools such as artificial insemination and sexed semen (Weigel et al., 2017). More recently, the introduction of genomic tools for quantifying each animal’s production merit has also improved the genetic progress (Fleming et al., 2018). The improvement of management and environment have also contributed to a better production performance. These changes include optimization of feed and nutrition, monitoring of health, reproduction and welfare, better veterinary practices, housing and climate control and milking routines (Rajala-Schultz et al., 1999b; Bach et al., 2008; Balaine et al., 2020).

Milk production is crucial for a dairy farm’s profitability, and a substantial part of the economic challenge is linked to losses of milk yield and quality due to health issues (van Soest et al., 2016, 2019; Liang et al., 2017). These losses manifest in altered milk yield dynamics seen as perturbations in the lactation curve (Hertl et al., 2014; Ben Abdelkrim et al., 2019). Both systemic and local effects of disease and inflammation contribute to these perturbations. The systemic effects are mainly caused by e.g., a decreased feed intake or a reallocation of energy to the immune system fighting the infection (Daniel et al., 2018). Local effects, especially in the case of intramammary infections, can be due to damage of the blood-milk barrier in the udder caused by somatic cell inflow or pathogen toxins (Heikkilä et al., 2018).

In the past, milk yield performance and dynamics were mainly analyzed in the context of genetic studies using low-frequency test-day data, for example for national herd health and genetic improvement programs (ICAR, 2014). Although proven useful, these low-frequency measurements do not allow for characterizing specific perturbation dynamics such as development and recovery rates, which reflects whether and to what extent a cow returns to normal production levels after a challenge. This kind of information, however, is useful to determine a cow’s robustness and resilience, and to compare her response to that of herd mates subjected to similar challenges, e.g. during heat stress episodes.

With high-frequency milk meter data, advanced computation, and storage capacities becoming widely available, the milk yield dynamics can be studied in more detail than before. This opens opportunities for precision phenotyping and targeted monitoring, starting from actual insight in lactation and disease dynamics, and including the aforementioned recovery and development rates and production losses. In order to study perturbations, first the theoretical unperturbed lactation curve has to be estimated, for example as proposed by Adriaens et al. (2018), Ben Abdelkrim et al. (2019), Adriaens et al. (2020) and Poppe et al. (2020). The deviations from this unperturbed curve can then serve as the basis for detecting and characterizing the deviations in milk yield and identification of challenges, especially in absence of detailed and reliable health registers.

In this study, we hypothesized that studying the milk losses and dynamics of perturbations can help to gather insights in a) how a dairy cow reacts to physiological and environmental challenges characterizing specific development and recovery phases of the perturbations; and how milk yield dynamics during challenges vary with lactation stage and parity. To this end, we present a methodology to calculate and characterize specific milk yield dynamics in perturbations based on individual daily milk yield measurements (**DMY**). Using this approach, the frequency, timings, milk losses and development and recovery rates of perturbations in the lactation curve of dairy cows were studied. Additionally, we tested the differences between the mean values of the characteristics across lactation stage and parity groups.

## MATERIALS AND METHODS

### Data Collection, Selection and Preprocessing

Back-ups of the data management software of automatic milking systems (**AMS**) were collected at 30 dairy farms with a Lely AMS (Lely Industries N.V., Maassluis, the Netherlands) and 28 farms with a DeLaval AMS (DeLaval, Tumba, Sweden) throughout Belgium, the south of the Netherlands and England. Using Microsoft SQL Server Management Studio 18 (SMSS, Microsoft, Redmond, WA, USA), these back-up files were restored and the data tables containing the historical DMY data and cow and lactation identification were extracted. All further data processing was done in Matlab R2019b (The MathWorks Inc., Natick, MA, USA). An individual table was constructed for each farm containing unique cow and lactation identifiers including birth and calving dates and the DMY together with their corresponding measurement dates and days in milk (**DIM**).

From these tables, lactations were selected based on the following 2 criteria: (1) the DMY data were available from before DIM 5 to at least DIM 305; and (2) no more than 2 gaps of at most 5 days each were present in the lactation curves. Only the first 305 days of each lactation were included in the analysis for standardization purposes. This avoids bias in the analysis caused by last part of the lactation curves, which can be substantially influenced by the gestation stage, feed changes towards dry-off etc. (Dematawewa et al., 2007; Ben Abdelkrim et al., 2019).

The DMY data from the DeLaval software back-ups differed from the DMY data from the Lely software back-ups because the first takes the sum of the milkings of that day as the daily value, while the latter corrects for the varying number of milkings between days typical for AMS data. All other differences are considered negligible. To obtain a similar variability in the DMY across all farms, the DMY data from the Delaval farms were corrected by replacing each value with the average of the current and previous day.

### Unperturbed Lactation Curve

Because health and treatment records are often incomplete and unreliable (Stevens et al., 2016) and because not all challenges are noticed or treated, health and treatment registers were not considered in this study to mark perturbations. Moreover we did not want to limit the study to perturbations caused by clinical events. Alternatively, high-frequency and consistent milk yield measurements were used to obtain a general and robust methodology for the identification of challenges and perturbations. This allows for inclusion of all farms for which high-granularity time-series measurements are available. To this end, the deviations from an unperturbed lactation curves (**ULC**) were calculated. These ULC represent the assumed theoretical production dynamics in absence of perturbations.

To calculate each lactations’ ULC, we assumed that the DMY during the first 305 days after calving follow the theoretical shape of the incomplete gamma function, referred to as the Wood model (Wood, 1967, Eq. 1).

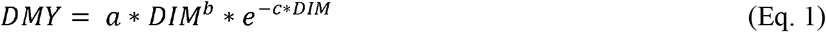

Wood’s model consists of 3 parameters, from which the *a* parameter mainly determines the scaling of the curve, whereas *b* and *c* specify the moment of peak production and slope. We preferred the Wood model over other lactation models because of its simplicity, its stability in the presence of missing data, the computational ease and its suitability for the purpose of this study (Adriaens et al., 2018; Ben Abdelkrim et al., 2019). To estimate the ULC that represents the milk production potential in absence of perturbations, an iterative fitting procedure was implemented to gradually remove milk yield data during perturbations. The methodology is visualized in Figure 1 and is as follows:

1. *In this first iteration (i = 1), fit the non-linear lactation curve model (**ULC_1_**, red dashed line in Figure 1) on all DMY data of the lactation (dark blue circles).*
2. *Calculate the residuals from this model, their standard deviation **SD_1_** and the root mean squared error (**RMSE_1_**);*
3. *Remove all DMY data below ULC_1_ −1.6*SD_1_ (red circles) to obtain the filtered DMY data resulting from the first iteration (i = 1). 1.6*SD was chosen as a threshold to exclude all data points outside the 95% CI;*
4. *In this next iteration (i), fit the non-linear lactation curve model (**ULC_i_**, light blue dashed line) on the filtered DMY data resulting from the previous iteration (i-1);*
5. *Calculate the residuals from this model, their standard deviation **SD_i_** and the root mean squared error (**RMSE_i_**);*
6. *Remove all DMY data below ULC_i_ – 1.6*SD_i_ (light blue circles) to obtain the filtered DMY data resulting from the current iteration (i);*
7. *Repeat steps 4 to 6 until the improvement in RMSE_i_ of the current iteration (i) compared to the previous iteration (RMSE_i-1_) is smaller than 0.1 kg or after 20 iterations (final model, **ULC_F_** depicted as the thick orange line).*

**Figure 1.**
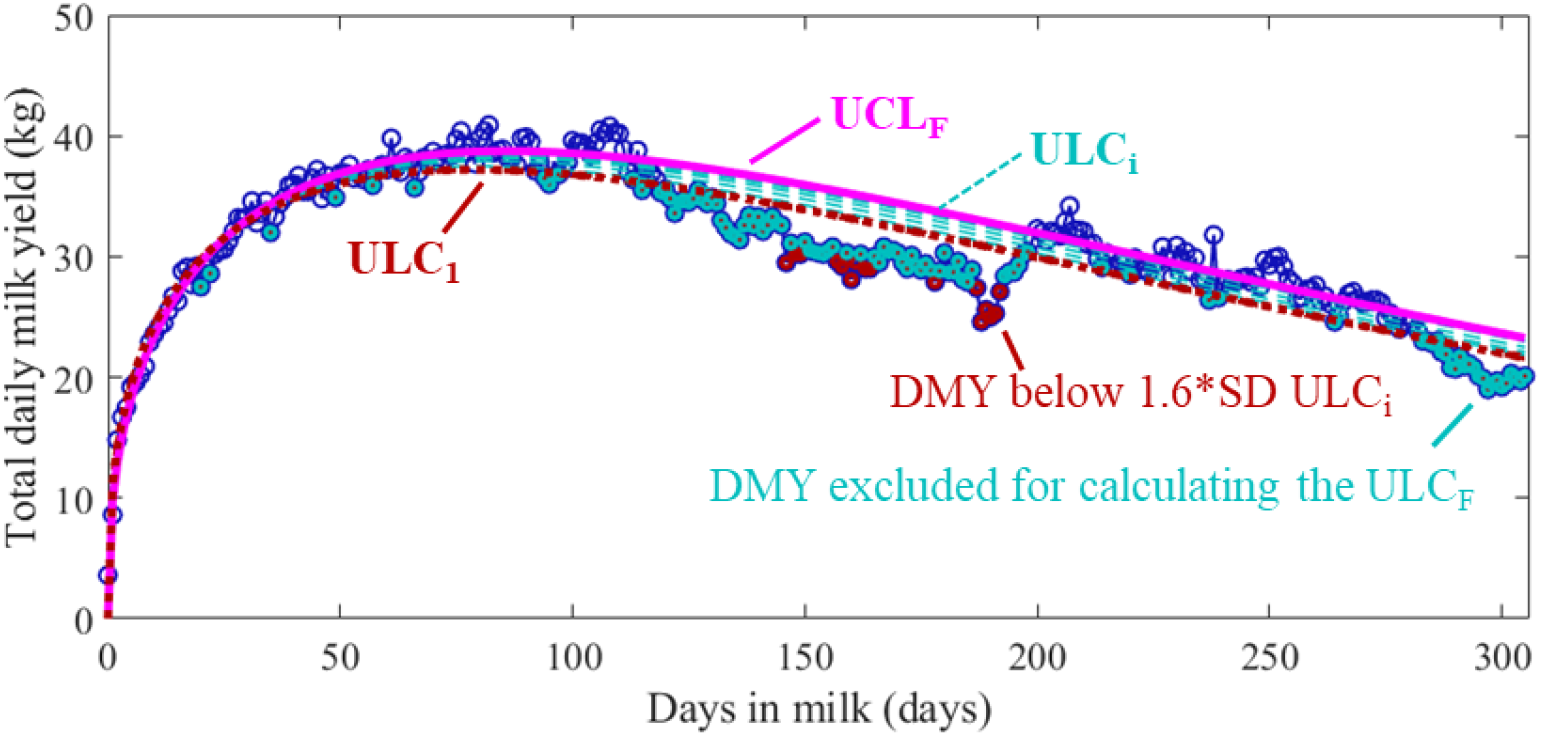
Example of the initial and final unperturbed lactation curves (ULC) obtained using an iterative fitting procedure that excludes perturbations. The red dotted line represents the initial ULC_1_, fitted on all daily milk yields (DMY). The red dots represent the milkings below this initial fit minus 1.6 * the standard deviation of the residuals, which are excluded during the first iteration. After 4 iterations, the magenta curve (ULC_F_) was obtained, fitted on all the blue data and thus, excluding the DMY in perturbations indicated in red and cyan.

For each iteration, the parameters (*a, b, c)* of the previous iteration are used as the initial values for the next iteration. The non-linear fitting procedure used the trust-region-reflective algorithm with at most 100 iterations and a lower boundary of 0 on the parameters to find the optimal solution. The resulting ULC_F_ are taken as the reference representing the total DMY that would have been achieved in absence of perturbations. Persistency of these ULC_F_ was calculated as the decrease in predicted DMY between DIM 205 and 305 expressed in kg.

### Detection of Perturbations

To detect perturbations, the ULC_F_ of each lactation was subtracted from the DMY to obtain the residuals, which are the deviations from the ULC_F_. These residuals are expected to vary around 0 in the absence of a perturbation, while during perturbations they will be consistently negative. A perturbation was defined as a period of at least 5 successive days of negative residuals for which the DMY dropped at least once below 80% of the expected yield (ULC_F_) as also discussed in Adriaens et al. (2020). The start and end DIM of each perturbation were marked by respectively the first and last residual below 0.

### Characterization of Perturbations

The DMY of each perturbation was smoothed using a third order Savitsky-Golay polynomial smoother. This unweighted least squares method filters out the effect of e.g. autocorrelation between days, for example when the milking process was not entirely completed resulting in a low DMY on one day and a high DMY the next day. Based on the minimum of the resulting smoothed values, the timing of the maximal loss (**TML**, i.e., the ‘center’ = day 0 of the perturbation) of each perturbation was determined. In the rest of the analysis, the days before the TML were considered as the development phase, while the days after the TML represent the recovery phase. An example of the raw and the smoothed data for one perturbation is shown in Figure 2. The red square represents the maximal loss, and the vertical dashed line indicates the TML.

**Figure 2.**
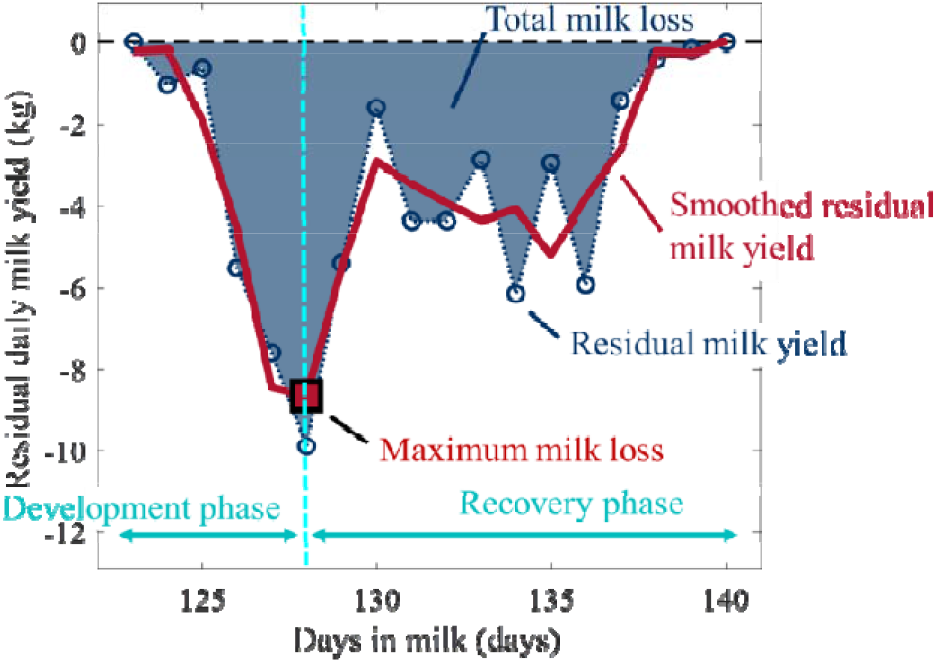
Example of a perturbation, its maximum milk loss and the resulting development and recovery phase. The blue circles indicate the residuals calculated using the unperturbed lactation curve. The red thick line represents the smoothed residuals from which the minimum serves as the reference to determine the development and recovery phase. The milk loss is calculated as the sum of the residuals.

### Statistical Analysis

For each perturbation, a number of characteristics were calculated including 1) the total duration and the duration of the development and recovery phase; 2) the slopes of both phases as the average difference between the smoothed DMY per day (in kg/day); and 3) the milk losses. These milk losses were calculated as the sum of the non-smoothed residuals of each perturbation, in total as well as for the development and recovery phases separately, and expressed both in absolute values (kg milk) and relative losses (% from the predicted milk yield). For in total 14 characteristics, the mean values were compared in a 2-way ANOVA with 2 groups (parity and lactation stage) and their interaction. The null hypothesis for this analysis was: “the mean of each characteristic in each parity and lactation stage group are the same”. The parity factor had 3 levels (first, second or third and higher), while for the lactation stage 4 levels were used (early, ‘E’ = DIM 0 to 63, mid-early, ‘ME’ = DIM 64 to 138, mid-late, ‘ME’, = 139 to 216, late, ‘L’ = 217 to 305). The split for lactation stage was based on the number of perturbations detected to obtain an approximately balanced amount of perturbations per lactations stage and parity group. Moreover, this split nicely represents the different phases of the lactation curve dynamics in the first 305 days after calving, where the ‘early’ level will reflect the increasing part, the ‘mid-early’ level will contain the peak, and ‘mid-late’ and ‘late’ the approximately linear part of most of the lactation curves. If the ANOVA pointed out significant effects of the factors, a Bonferroni mean comparison was carried out to determine significant differences between the means of the groups. Bonferroni was preferred over Tukey HSD to avoid capitalization of chance on Type I statistical errors when a high amount of means is compared. In this study this potentially summed up to 66 comparisons, testing the differences in means for 12 combinations of lactation stage (No. groups = 4) and parity (No. groups = 3).

## RESULTS

### Data Set Description

From the data set of 58 farms (30 DeLaval, 28 Lely), no cows were selected for 8 farms because they either had a high amount of gaps (missing data, e.g. because of technical or data storage errors) or did they did not cover a long enough time period (e.g. too recent installation of the AMS system or a reset of the historical data after upgrade to a new software version of the robot). The other 50 farms had at least 1 cow for which 305 days of uninterrupted data were available. Thirty of these farms milked with Lely, and 20 with Delaval AMS. The resulting data set consisted of 9,681 unique cows and 16,640 unique lactations from which respectively 5,590, 4,543 and 6,507 from the first, second and third or higher parity. Only the data of the first 305 days (DIM 0 to 305) were considered, resulting in a total data set size of more than 50 million DMY records. The summary statistics of the selected data set are given in Table 1, and Figure 3 shows the average and average ±3*SD for the lactation curve data together with 100 randomly selected individual lactation examples per parity (first, second and third or higher). The random selection was done by sampling 100 numbers from a uniform distribution.

**Figure 3.**
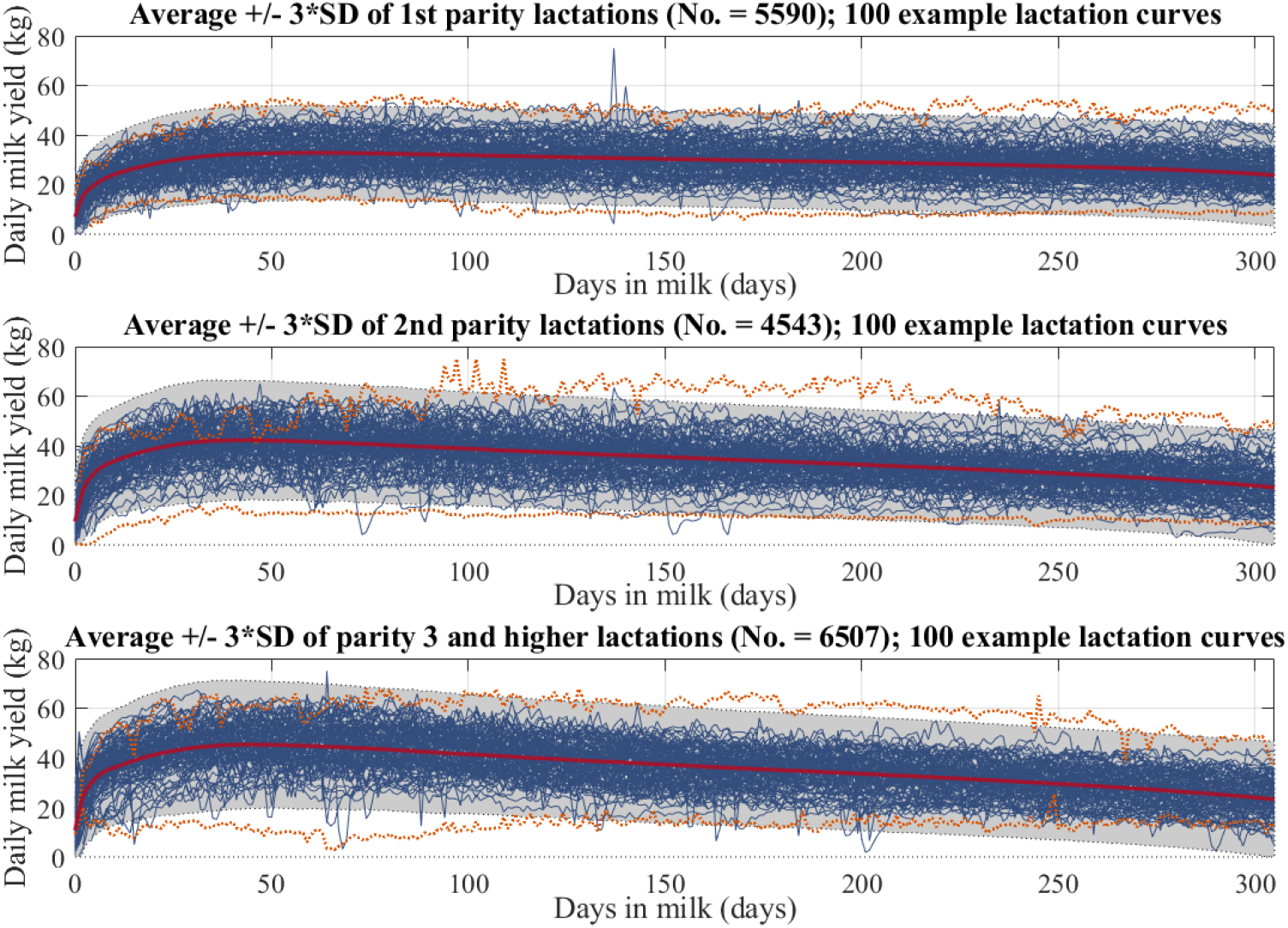
Overview of the raw daily milk yield (DMY) data for each parity (first, second and third or higher) over time. The gray background shows the mean ± 3* standard deviation (SD), the orange dotted lines show the curves with respectively the highest and lowest milk yield summed over the first 305 days. The blue thin lines show 100 randomly chosen examples per parity, while the thick red line gives the average lactation curve for that parity.

**Adriaens, Table 1.** Summary statistics of the selected data set

| | No. for the entire data set | Mean ± SD<sup>2</sup> over farms | Range over farms [minimum ; maximum] |
| --- | --- | --- | --- |
| **No. farms included** | 50 |  |  |
| Lely AMS<sup>1</sup> | 30 |  |  |
| Delaval AMS<sup>1</sup> | 20 |  |  |
| **Time period covered** [years] | | 6.6 ± 3.4 | [0.8 ; 14.0 ] |
| **No. of unique cows** | 9681 | 193.6 ± 139.4 | [ 1 ; 879 ] |
| **No. of unique lactations** | 16640 | 332.8 ± 299.7 | [ 1 ; 1968 ] |
| Parity 1 | 5590 | 111.8 ± 105.6 | [ 0 ; 701 ] |
| Parity 2 | 4543 | 90.9 ± 85.4 | [ 0 ; 565 ] |
| Parity 3+ | 6507 | 130.1 ± 115.2 | [ 0 ; 702 ] |
| **Average age at first calving** [years] | | 2.1 ± 0.1 | [ 1.9 ; 2.4 ] |
| **Average daily milk yield in first 305 days** [kg] |  |  |  |
| Parity 1 | | 29.0 ± 3.2 | [ 17.5 ; 34.2 ] |
| Parity 2 | | 34.1 ± 3.7 | [ 21.5 ; 41.6 ] |
| Parity 3+ | | 35.6 ± 3.6 | [ 25.6 ; 42.2 ] |
| **Total sum of milk yield in first 305 days** [kg] |  |  |  |
| Parity 1 | | 8841.4 ± 974.9 | [ 5296.3 ; 10466.5 ] |
| Parity 2 | | 10406.2 ± 1122.5 | [ 6529.7 ; 12692.3 ] |
| Parity 3+ | | 10885.7 ± 1120.6 | [ 7770.2 ; 12848.8 ] |

<sup>1</sup>AMS: automated milking system; <sup>2</sup>SD: standard deviation

### Unperturbed Lactation Curves

An overview of the summary statistics of the 16,640 lactations is given in Table 2. The procedure to estimate the unperturbed curve converged on average in 3.7 ± 1.5 (mean ± SD) iterations, and resulted in an average RMSE of 3.6 ± 1.4 kg. The RMSE, which is a measure for goodness-of-fit and the average deviation of all daily milk yield data from the estimated unperturbed curve, increased with parity. This can be due to the lower milk production in general for primiparous cows, but can also be due to a higher number or severity of perturbations. As expected, the scaling parameter of the Wood models representing the ULC_F_ increased with parity from 15.7 ± 4.6 for first parity lactations to 23.7 ± 6.5 for third and higher lactations. This translates in an average peak yield of 33.9 ± 5.7, 42.8 ± 7.2 and 46.1 ± 7.3 kg for first, second and third and higher lactations respectively. For first parity lactations, this peak was estimated at DIM 79.5 ± 36.5, while it was respectively at DIM 53.3 ± 20.2 and 50.9 ± 18.3 for second and third and higher parity lactations. The SD for peak DIM was clearly higher for primiparous than for multiparous cows. This is due to the generally flatter (i.e., more persistent) shape of the lactation curve in the first parity, making the estimation of the exact peak less robust. This was confirmed by the average persistency, expressed as the decrease in kg milk from DIM 205 to DIM 305. The DMY for first parity cows decreased with on average 4.1 ± 2.2 kg, while second and third and higher parity cows decreased with respectively 7.8 ± 2.7 and 9.1 ± 2.9 kg. Similar conclusions can be drawn from Figure 4, in which for each parity 50 randomly chosen curves are plotted together with the average, minimum and maximum estimated ULC_F_. The red dots represent the peak yield, while the orange dotted lines show the mean ± 3*SD of the ULC_F_ for each parity. The grey shading represents the mean ± 3*SD of the raw total DMY data. The upper orange line is close to the upper limit of the grey shading, as can be expected because only data belonging to perturbations (i.e., strongly negative deviations) were deleted in the iterative procedure to obtain the ULC_F_.

**Figure 4.**
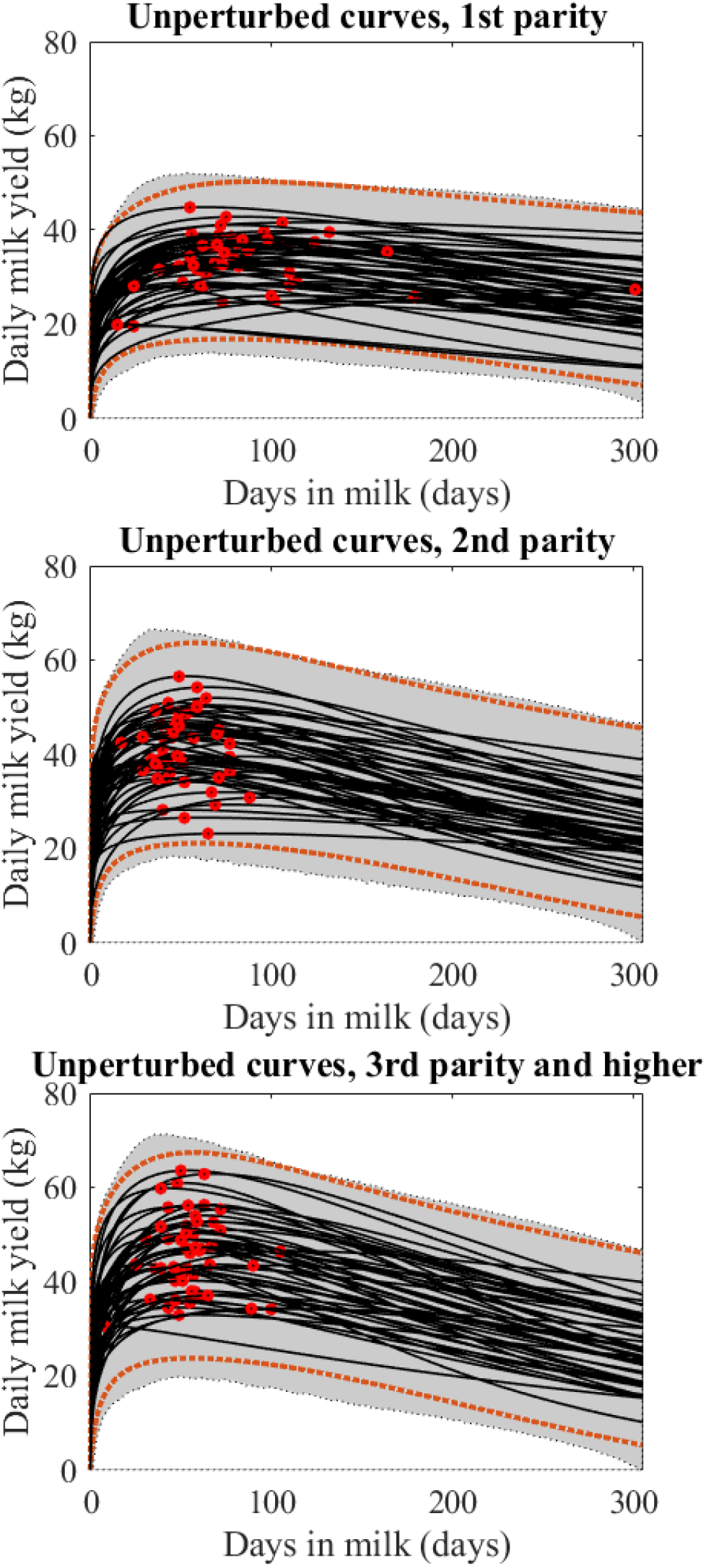
Fifty randomly chosen examples of the final unperturbed lactation curves (ULC_F_) for each parity. The red dots represent the peak yield, while the orange dotted line gives the mean ± 3 * standard deviation of these UCL_F_. The grey shading represents mean ± 3*SD of the raw daily milk yield data.

**Adriaens, Table 2.** Summary statistics of the unperturbed lactation curves

MILK YIELD DYNAMICS DURING PERTUBATIONS Adriaens, Table 2. Summary statistics of the unperturbed lactation curves
| | All<br>(mean ± SD<sup>4</sup>) | 1<sup>st</sup> parity<br>(mean ± SD<sup>4</sup>) | 2<sup>nd</sup> parity<br>(mean ± SD<sup>4</sup>) | 3<sup>rd</sup> parity and higher<br>(mean ± SD<sup>4</sup>) |
| --- | --- | --- | --- | --- |
| **Wood parameters** |  |  |  |  |
| a | 20.6 ± 6.9 | 15.7 ± 4.6 | 22.4 ± 6.3 | 23.7 ± 6.5 |
| b | 0.23 ± 0.09 | 0.23 ± 0.09 | 0.22 ± 0.08 | 0.23 ± 0.09 |
| c | 0.0035 ± 0.0014 | 0.0027 ± 0.0011 | 0.0037 ± 0.0013 | 0.0040 ± 0.0014 |
| No. of iterations<sup>1</sup> | 3.6 ± 1.3 | 3.3 ± 1.3 | 3.5 ± 1.4 | 3.8 ± 1.6 |
| RMSE<sup>2</sup> [kg] | 3.6 ± 1.4 | 3.0 ± 1.0 | 3.6 ± 1.3 | 4.0 ± 1.4 |
| **Lactation curve characteristics** |  |  |  |  |
| Peak yield [kg] | 41.1 ± 8.6 | 33.9 ± 5.7 | 42.8 ± 7.2 | 46.1 ± 7.3 |
| Peak DIM<sup>3</sup> [days] | 61.1 ± 29.3 | 79.5 ± 36.5 | 53.3 ± 20.2 | 50.9 ± 18.3 |
| Persistency [kg/100 days] | 7.0 ± 3.4 | 4.1 ± 2.2 | 7.8 ± 2.7 | 9.1 ± 2.9 |

<sup>1</sup> No. of iterations: Number of iterations needed to establish the final unperturbed lactation curve (ULC<sub>F</sub>); <sup>2</sup> RMSE: root mean squared error of the final unperturbed curve; <sup>3</sup> DIM: days in milk; <sup>4</sup> SD: standard deviation

### Number and Timing of Perturbations

The data set contained a total of 62,406 perturbations, defined as periods of at least 5 successive days of milk production below the ULC_F_ from which one day less than 80% of the expected DMP. From these perturbation, respectively 21,442, 16,499 and 24,465 belonged to first, second and third or higher parity lactations. This corresponds to 3.7 ± 1.8 perturbations per lactation on average. Only 172 lactations (1.0 %) had no perturbations, while respectively 7.3, 19.1, 22.7, 19.2, 13.8 and 16.8 % of the lactations had 1, 2, 3, 4, 5 or 6 or more perturbations. The average duration of each perturbation was 19.7 ± 20.6 days. For 44.6% of the perturbations the duration was 10 days or shorter, while 37.7% of the perturbations lasted between 10 and 30 days. The remainder of the perturbations lasted more than 30 days.

The timing of the perturbations is graphically represented in Figure 5, for each 10-day time period of the lactation showing the proportion of perturbations that start in that period. In total, 13.3% of all perturbations started in the first 10 days of lactation, and this percentage was even higher for first parity cows (15.8%). Compared to the first 10 DIM, less perturbations were detected between DIM 10 and 60, which might be because new perturbation can only be detected when the previous perturbation was completely over. Also, between day 60 and 120 and after day 250, a clear increase in number of perturbations was detected.

**Figure 5.**
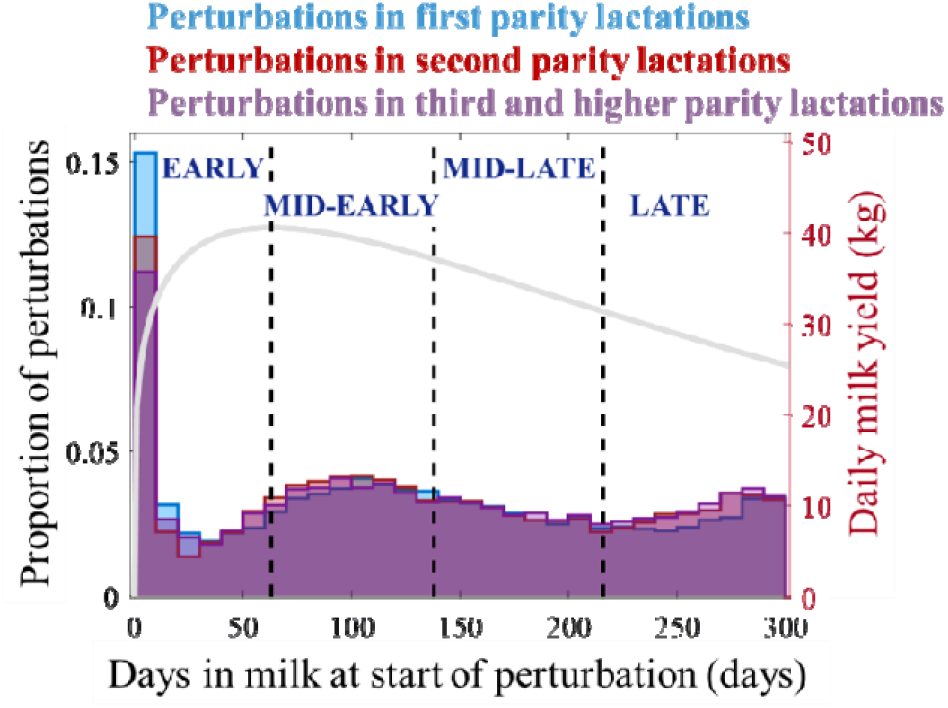
Proportion of the start of the perturbations across lactation stage for each parity. Based on the number of perturbations, lactation stage groups were determined as ‘early’ = 0 to 63 DIM, ‘mid-early’ = 64 to 138 DIM, ‘mid-late’ = 139 to 216 DIM and ‘late’ = 217 to 305 DIM. The grey thick line shows the average unperturbed lactation model for all parities.

A further overview of summary statistics for number, duration and timing of the perturbations is given in Table 3. The percentage lactations without recovery are the lactation from which the last perturbation did not get back to the ULC_F_, thus the residuals remained negative, up till at least DIM 305.

**Adriaens, Table 3.**
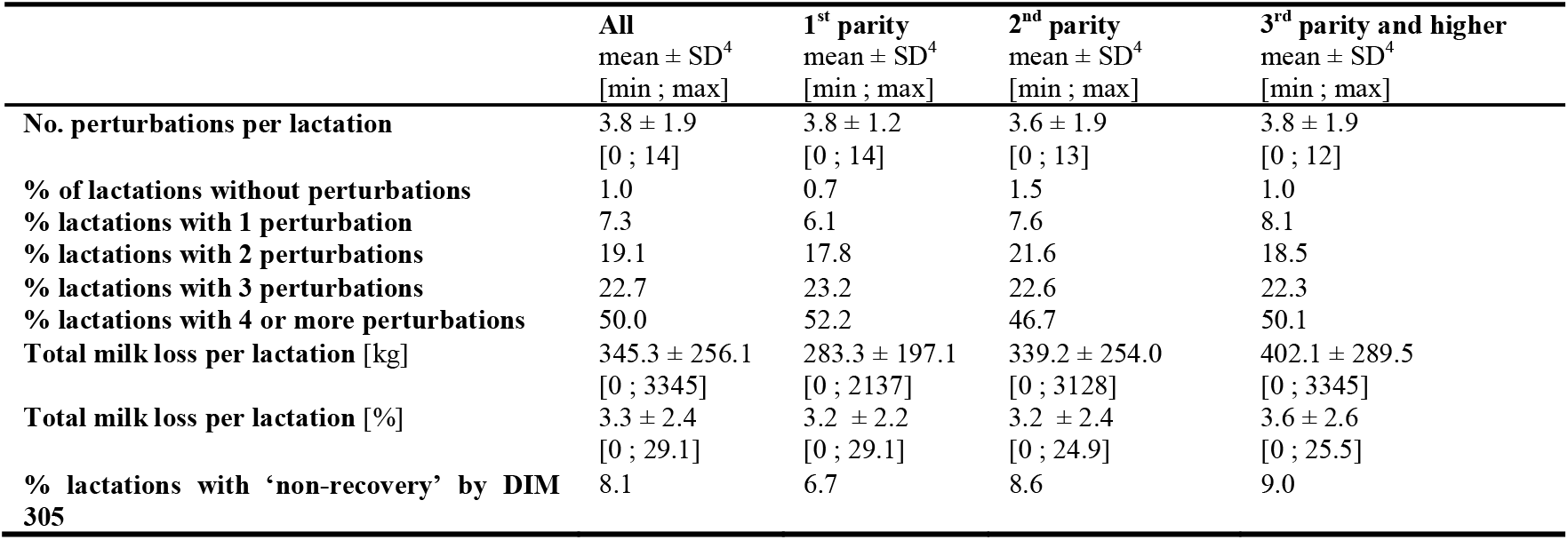
Overview of the perturbations’ number, timing and milk losses per lactation (305 days)

### Perturbation Characteristics

Table 4 shows the perturbation characteristics overall and per parity. On average, 92.1 kg milk was lost per perturbation, with a very high standard deviation of 135 kg. Perturbations in higher parity cows have on average higher losses than first parity perturbations. Also when expressed in kg per day, milk losses increase with parity.

**Adriaens, Table 4.** Summary statistics of perturbations overall and per parity

MILK YIELD DYNAMICS DURING PERTUBATIONS Adriaens, Table 4. Summary statistics of perturbations overall and per parity
| | All<br>mean ± SD<sup>1</sup><br>[min ; max] | 1<sup>st</sup> parity<br>mean ± SD<br>[min ; max] | 2<sup>nd</sup> parity<br>mean ± SD<br>[min ; max] | 3<sup>rd</sup> parity and higher<br>mean ± SD<br>[min ; max] |
| --- | --- | --- | --- | --- |
| **Total milk loss [kg]** | 92.1 ± 135.8<br>[4.6 ; 3345] | 73.8 ± 104<br>[4.6 ; 1956] | 93.4 ± 136<br>[5.3 ; 3029] | 107.2 ± 156.5<br>[5.1 ; 3345] |
| **Average milk loss per day [kg/day]** | 3.5 ± 1.8<br>[0.5 ; 30.6] | 3.0 ± 1.4<br>[0.6 ; 15.2] | 3.6 ± 1.8<br>[0.6 ; 20.0] | 3.9 ± 2.0<br>[0.6 ; 30.6] |
| **Duration [days]** | 19.8 ± 20.7<br>[5 ; 206] | 18.8 ± 20.0<br>[5 ; 186] | 19.8 ± 20.9<br>[5 ; 179] | 20.5 ± 21.0<br>[5 ; 206] |
| **Development phase** |  |  |  |  |
| **slope [kg/day]** | -2.3 ± 3.0<br>[-41.3 ; 0.11] | -2.0 ± 2.4<br>[-28.0 ; 0.08] | -2.5 ± 3.2<br>[-41.3 ; 0.09] | -2.6 ± 3.4<br>[-40.8 ; 0.11] |
| **duration [days]** | 10.1 ± 12.6<br>[1 ; 142] | 9.3 ± 11.5<br>[1 ; 132] | 10.3 ± 13.0<br>[1 ; 132] | 10.6 ± 13.3<br>[1 ; 142] |
| **milk loss [kg]** | 48.2 ± 78.7<br>[0.9 ; 1849] | 38.0 ± 58.0<br>[0.9 ; 1491] | 49.7 ± 81.3<br>[2.0 ; 1573] | 56.0 ± 90.8<br>[1.18 ; 1849] |
| **Recovery phase** |  |  |  |  |
| **slope [kg/day]** | 1.5 ± 1.8<br>[0.02 ; 30.5] | 1.3 ± 1.53<br>[0.03 ; 22.4] | 1.6 ± 1.9<br>[0.04 ; 22.4] | 1.6 ± 1.9<br>[0.02 ; 30.5] |
| **duration [days]** | 11.6 ± 15.1<br>[0 ; 166] | 11.2 ± 14.6<br>[0 ; 141] | 11.6 ± 15.2<br>[0 ; 162] | 12.0 ± 15.5<br>[0 ; 166] |
| **milk loss [kg]** | 47.8 ± 94.7<br>[0 ; 2713] | 38.4 ± 71.8<br>[0 ; 1883] | 47.8 ± 94.8<br>[0 ; 2712] | 56.2 ± 110<br>[0 ; 2273] |
| **% perturbations of duration**<br>**(cumulative)** |  |  |  |  |
| **$\leq 10$ days** | 44.6 | 46.9 | 45.2 | 42.3 |
| **$\leq 15$ days** | 61.2 | 64.0 | 61.1 | 58.9 |
| **$\leq 20$ days** | 71.2 | 73.7 | 70.9 | 69.3 |
| **$\leq 25$ days** | 77.6 | 79.7 | 77.2 | 75.6 |
| **$\leq 30$ days** | 82.2 | 83.8 | 82.0 | 80.8 |
| **$> 30$ days** | 17.8 | 16.2 | 18.0 | 19.2 |

<sup>1</sup> SD: standard deviation.

In 57.2% of the perturbations, the development slope was steeper than the slope in the recovery phase, with respectively −2.3 kg and +1.5 kg per day. On average, the development phase was also 1.5 days shorter than the recovery phase, the latter lasting for on average 11.6 days. Milk losses in both phases seem to be comparable. In absolute numbers, perturbations of the first parity are less severe (less milk losses, slower development and recovery rates) than perturbations in higher parity cows. Because we decided to cut off the lactations at 305 DIM, no recovery was reached for some perturbations at the end of the lactation. When leaving out these ‘non-recovery’ perturbations, the duration of the development phase decreased to 8.3 ± 10.3 days. The duration of these perturbations was on average 29 ± 19 days, which is 10 days longer than the average duration of all perturbations together (19.8 ± 20.7 days). Of all perturbations, 44.6 % lasted 10 days or shorter, while respectively 61.2, 71.2, 77.6 and 82.2% lasted shorter than 15, 20, 25 or 30 days. The other 17.8% had a duration of more than 30 days.

### Statistical Analysis

Grouping of the data for parity and lactation stage resulted in the following data split (Table 5), and the difference between parity and lactation stage were tested for 14 different characteristics. Seven of these characteristics are linked to milk losses, while the other 7 are related to the shape and dimension of the perturbations. Their names, units, definitions, the range of the means of the groups and the *P*-values of the ANOVA are given in Table 6. The degrees of freedom of each test were respectively 2, 3 and 6 for the parity, lactation stage and interaction between both factors, and the tests had 57,333 and 62,394 error degrees of freedom for respectively the characteristics involving specifically the recovery phase and the other characteristics.

**Adriaens, Table 5.**
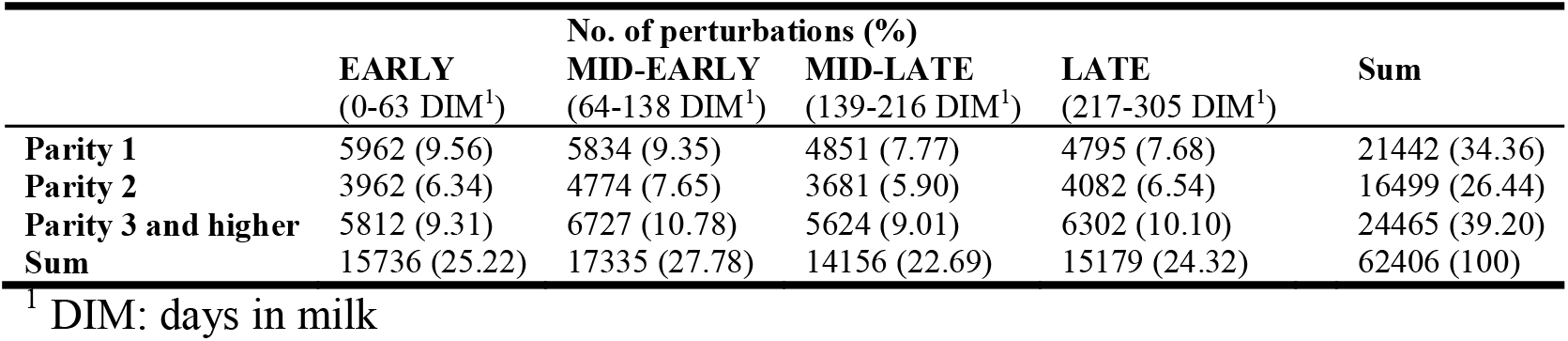
Number of perturbations in each combination of lactation stage and parity group

|  | No. of perturbations (%) |  |  |  | Sum |
| --- | --- | --- | --- | --- | --- |
|  | EARLY<br>(0-63 DIM<sup>1</sup>) | MID-EARLY<br>(64-138 DIM<sup>1</sup>) | MID-LATE<br>(139-216 DIM<sup>1</sup>) | LATE<br>(217-305 DIM<sup>1</sup>) |  |
| Parity 1 | 5962 (9.56) | 5834 (9.35) | 4851 (7.77) | 4795 (7.68) | 21442 (34.36) |
| Parity 2 | 3962 (6.34) | 4774 (7.65) | 3681 (5.90) | 4082 (6.54) | 16499 (26.44) |
| Parity 3 and higher | 5812 (9.31) | 6727 (10.78) | 5624 (9.01) | 6302 (10.10) | 24465 (39.20) |
| Sum | 15736 (25.22) | 17335 (27.78) | 14156 (22.69) | 15179 (24.32) | 62406 (100) |

<sup>1</sup> DIM: days in milk

**Adriaens, Table 6.** Names, units and description of the characteristics of the individual perturbations for which the effect of parity and lactation stage were tested in a 2-way ANOVA. The range of the absolute value of the means for each group (parity*lactation stage) and respective P-values are also given.

| Characteristic | Unit | Definition | Range groups | P-value ANOVA |  |  |
| --- | --- | --- | --- | --- | --- | --- |
|  |  |  | [min ; max] | Parity | LS<sup>1</sup> | Parity*LS |
| MILK LOSSES |  |  |  |  |  |  |
| LOSS-DAY | kg/day | Average loss per day | [2.9 ; 4.4] | <0.0001 | <0.0001 | <0.0001 |
| LOSS | kg | Sum of the residuals per perturbation (DMY-ULC<sub>F</sub>) | [58.1 ; 140.3] | <0.0001 | <0.0001 | <0.0001 |
| LOSS-DEV | kg | Sum of the residuals in the development phase per perturbation | [26.4 ; 73.3] | <0.0001 | <0.0001 | <0.0001 |
| LOSS-REC<sup>2</sup> | kg | Sum of the residuals in the recovery phase per perturbation | [21.2 ; 80.2] | <0.0001 | <0.0001 | <0.0001 |
| RELLOSS | %/day | Average loss per day in % | [11.1 ; 15.0] | 0.0001 | <0.0001 | <0.0001 |
| RELLOSS-DEV | %/day | Average loss per day in % in the development phase | [14.0 ; 24.7] | <0.0001 | <0.0001 | <0.0001 |
| RELLOSS-REC | %/day | Average loss per day in % in the recovery phase | [10.6 ; 16.6] | <0.0001 | <0.0001 | <0.0001 |
| DIMENSIONS |  |  |  |  |  |  |
| DUR | days | Duration of perturbation | [13.6 ; 27.0] | <0.0001 | <0.0001 | <0.0001 |
| DUR-DEV | days | Duration of development phase | [5.6 ; 14.4] | <0.0001 | <0.0001 | <0.0001 |
| DUR-REC<sup>2</sup> | days | Duration of recovery phase | [6.9 ; 16.3] | <0.0001 | <0.0001 | <0.0001 |
| MML | kg | Minimum of smoothed perturbation | [7.2 ; 11.2] | <0.0001 | <0.0001 | <0.0001 |
| RELMML | % | % loss during minimum of smoothed perturbation | [24.3 ; 31.2] | <0.0001 | <0.0001 | <0.0001 |
| SLOPE-DEV | kg/day | Average slope during development phase (absolute value) | [1.3 ; 4.2] | <0.0001 | <0.0001 | <0.0001 |
| SLOPE-REC<sup>2</sup> | kg/day | Average slope during recovery phase | [1.3 ; 1.7] | <0.0001 | 0.0346 | <0.0001 |

<sup>1</sup> LS: lactation stage. <sup>2</sup> The perturbation characteristics for which less observations were available because no recovery was reached are indicated with the superscript 2. The ANOVA error degrees of freedom of these characteristics were 57,333 instead of 62,394.

As shown in Table 6, the effects of parity, lactation stage and their interactions were significant to highly significant for all 14 characteristics. The Bonferroni mean comparison tests pointed out which mean values and which trends were detected across parities and lactation stages. Moreover, for each characteristic the means of each group are graphically represented in Figure 6. Here, a darker color reflects a higher value and the greyscale is proportional to the mean values, such that similar colors indicate values closer to each other. The range of the means of each parity and lactation stage group is given at the right side of the figure, and E, ME, ML, L stand for ‘early’, ‘mid-early’, ‘mid-late’ and ‘late’ respectively.

**Figure 6.**
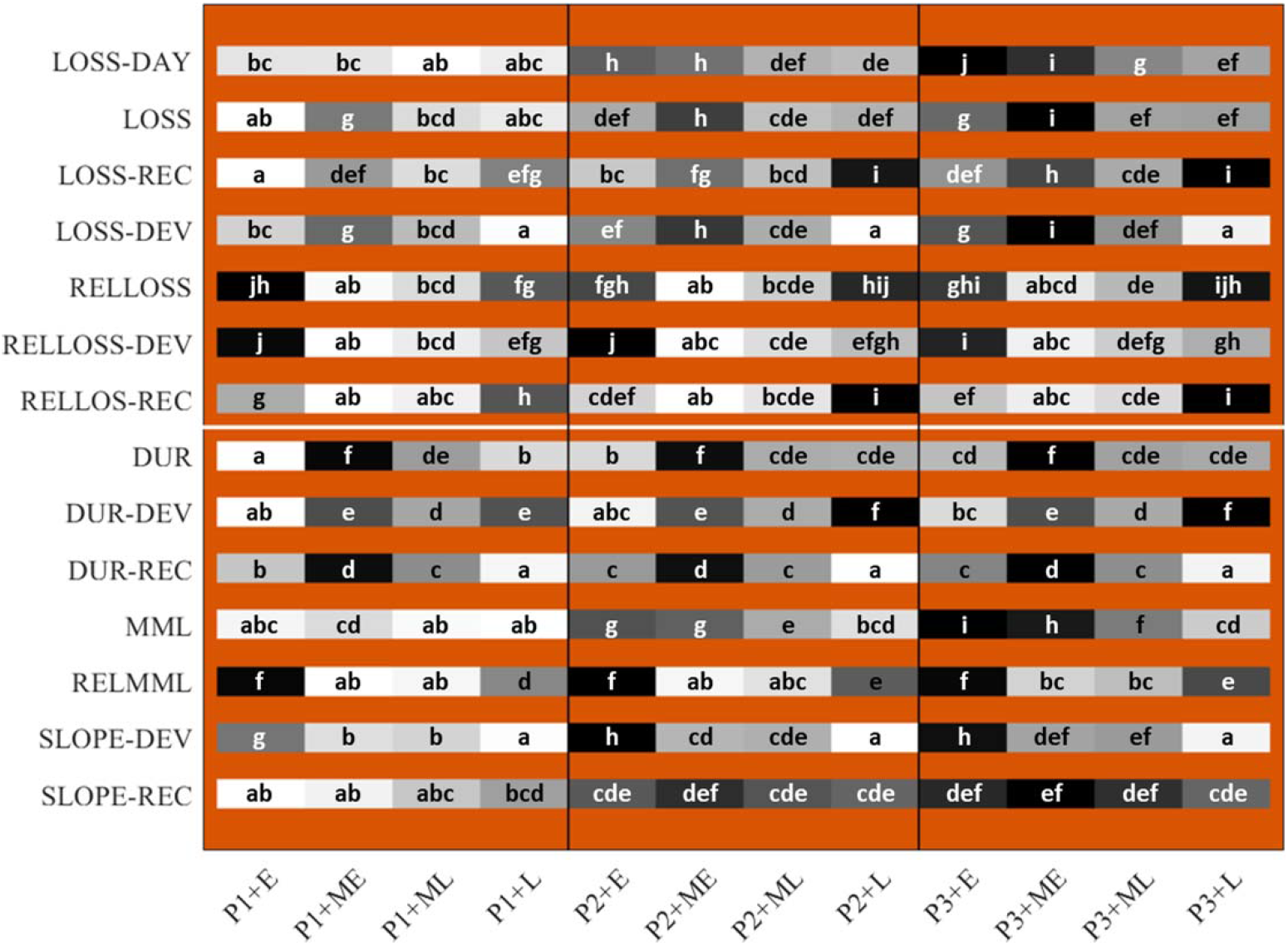
Graphical representation of the mean values for each perturbation characteristic. Higher values have a darker color and a letter code further in the alphabet, and the greyscale is proportional to the value such that more similar hues represent means closer to each other. Groups with the same letter code were not statistically different from each other. P indicates the parity (P1 = first parity, P2 = second parity, P3 = third or higher parity), and E, ME, ML and L represent the lactation stage classes ‘early’ = 0 to 63 days in milk, DIM, ‘mid-early’ = 64 to 138 DIM, ‘mid-late’ = 139 to 216 DIM and ‘late’ = 217 to 305 DIM respectively.

As expected, the milk losses per perturbation expressed in kg increase with parity, and are higher during peak lactation (LS = mid-early), probably because the expected milk yield is highest in this lactation stage. For the recovery phase, the milk losses per perturbation are lower at the end of the lactation, which might be due to the fact that the lactations were cut off at 305 DIM and because perturbations that did not recover were not included for the calculation of this characteristic. When considering relative losses expressed in % loss per day, early and late lactation perturbations clearly show a higher loss for each parity, likely because milk production is generally lower in these stages. Moreover, the high relative milk losses in early lactation are mainly related to the development phase, while those in late lactation are mainly linked to the recovery phase. Mid-early (so during peak production) perturbations last longest, which might be explained by the fact that cows in this lactation stage are under high metabolic stress. Therefore, health problems can have a stronger effect on the milk production dynamics and cows might experience a slower recovery. Indeed, the recovery phases are longer in mid-early lactation for each parity, while the development phases have the longest duration for the late-lactation perturbations. The latter can reflect the effect of chronic diseases in this stage, which is also supported by the low development slope. Other explanations for this are the possible change in diet towards dry-off and increased energy consumption of the fetus in pregnant animals. The slope of the development phase expressed in kg/day was found to decrease with parity. In early lactation, the slope of the development phase is clearly more steep for all parities, indicating sudden and clinical health problems (e.g. transition diseases). For the recovery slope, we can see that there is a clear parity effect, and first parity animals have a slower recovery than higher parity animals. Also, the slope of the recovery phase for first parity cows increases with lactation stage, while for second and third or higher parity animals the slope is higher in mid-early lactation. Additionally when considering the maximal milk loss of each perturbation expressed in % of the expected milk production (RELMML), early lactation perturbations tend to have higher milk losses compared to perturbations in the later lactation stages, again supporting the idea of more severe challenges in early lactation.

## DISCUSSION

### Lactation Modeling and Milk Losses

In this study we used the theoretical shape of the lactation curve (ULC) of AMS-milked dairy cows in the first 305 days to identify and characterize perturbations in the daily milk yield dynamics. To this end, we implemented an iterative, but computationally highly efficient method to estimate the presumed unperturbed lactation curve using the gamma function with three parameters (Wood model), as also proposed in Adriaens et al. (2018, 2020) and Ben Abdelkrim et al. (2019). Several other methods were tested for this study, including a 4^th^ order quartile regression model as proposed by Poppe et al. (2020) and the Dijkstra lactation model (Dijkstra et al., 1997). These methods showed very similar results which are therefore not presented in this manuscript. The Wood model is shown to describe dairy cows’ lactation curves in the first 305 days well, and only after the first 305 days and depending on the parity, more complex models perform better and are preferred (Dematawewa et al., 2007). While in the absence of severe health events most lactation curves show a typical growth towards a peak followed by a linear decrease, the very last part of the lactation often has more variation, strongly dependent on the lactation curve persistency, gestation stage and feeding status towards dry-off of the cows (Ehrlich, 2013). Accordingly, deviations from this linear decrease are often not only linked to health events, which was the interest of this study, but can also be attributed to a changing energy allocation upon fetus growth or changes in the animals’ diet to facilitate drying-off or to avoid obese animals in the dry period. Therefore, only the first 305 days of each lactation were taken into account for the analysis. Another asset for our choice for the Wood model besides the model fit and the computational advantage was its shape stability in the presence of missing data. This is crucial for the iterative procedure we used.

Negative deviations in milk yield for at least 5 days with at least one day below 80% of the expected yield were considered as perturbations and included in this study. In this way, short deviations caused by for example hardware or measurement errors (Bach and Busto, 2005), or linked to estrus (Gaillard et al., 2016) were excluded from the analysis. The idea was to focus on perturbations caused principally by clinical and subclinical health events and challenges, however not exclusively because the exact causes were not known.

Unfortunately, complete information on the actual health status of the animals was not available or reliable for all farms, and so we could not always discriminate between different perturbation’s causes. Data quality and reliability remains a challenge when using on-farm data (Hermans et al., 2017; Hudson et al., 2018). Besides health events, also changes in feed quality, social group changes or for example episodes of heat stress might lie at the basis of the detected perturbations (Dewhurst et al., 2002; Delaby et al., 2009; Bernabucci et al., 2014; Herve et al., 2019). For example towards the end of the lactation, these perturbations can be caused by change in diets towards dry-off and the energy consumption of the growing fetus. In these cases, milk yield was not expected to recover. On the other hand, when these perturbations are caused by health events, at least partial recovery would be expected. Studying specific milk yield dynamics, both at the end of the lactation or during specific health events, for example analyzed using research farm data for which reliable health and treatment registers are available, can be an interesting extension of the current study, but were outside the scope of the present analysis.

The effect of individual diseases on milk production in dairy cows is well described in literature, however often 1) using lower frequency data for example monthly test-day records or weekly yield aggregates (Rajala-Schultz et al., 1999b; Gröhn et al., 2004; Wilson et al., 2004; Hertl et al., 2014); or 2) averaging the milk losses from many cows with the same diseases together and comparing them with their healthy herd mates (Fourichon et al., 2000). For their thorough discussion, we refer to the work of Rajala-Schultz et al., (1999a; b), de Haas et al., (2002), Hand et al., (2012), and Hertl et al., (2014) amongst others. These studies confirm the present results in terms of order of magnitude of the daily milk losses, but did not describe specific dynamics or discriminate between development and recovery phases. In the present study, we also found that higher parities showed a larger number of perturbations, which can be due to higher incidence rates of clinical diseases (Lee and Kim, 2006). Similar to the findings of Carvalho et al., (2019), milk losses were higher for higher parity animals. When expressed in a relative way, these differences become minimal.

### Perturbation Characteristics

Only recently, large datasets of high-frequency milk yield time series became available, thanks to the installation of sensor technology and improved data storage capacities on farm. These datasets allow research on the individual lactation dynamics in more detail beyond averages or projected yield based on test days. Accordingly, also specific characteristics can be researched from these time series (Adriaens et al., 2020). The present study demonstrates a computationally efficient and intuitive method to characterize these perturbations post-hoc, i.e. after 305 days of lactation have been completed. Moreover, distinguishing between development and recovery phases facilitates studying the type of challenges to which cows are submitted and how they react on them. The methodology is not restricted to one specific type of perturbation. When this characterization is applied on a subset of perturbations caused by e.g. clinical mastitis, it can provide insights into the correlations between the perturbation characteristics and the infection, immune response, cure and treatment success. This allows not only for detection of a challenge, but also to look into the reaction in milk production on these challenges over time. When analyzing all the perturbations together, we noticed that the variability across perturbation characteristics is very high and that a ‘standard’ perturbation shape does not exist. As no clear correlations between the different characteristics were present, we did not report them for this study. This high variability might suggest that the perturbation characteristics can allow for the identification of differences between challenges and cows, to be used for example in monitoring or phenotyping applications (Dunne et al., 2018). Ultimately this would also allow for e.g. optimizing the parity composition on farm, taking into account trade-offs between production level, resiliency and robustness across parities.

We found an average of 3.8 perturbations per parity in the first 305 days corresponding to an average relative milk loss of 3.3% per lactation (range 0 to 29%, with 0 for cows without any perturbations detected). In dairy goat data, Ben Abdelkrim et al., (2019) identified 7.4 perturbations per lactation with losses between 2 and 19%. A major difference between our method and the method of Ben Abdelkrim et al., (2019) was that they did not exclude short perturbations below 5 days and that they considered also ‘perturbations within perturbations’, while we regarded each episode of negative residuals as a single perturbation. Accordingly, our results seem to confirm what was found by these authors in goat data.

The mean of each perturbation characteristic varied significantly with lactation stage and parity. In general, perturbations in early and mid-early lactation seem to be more severe with higher relative losses, faster development rates and slower recovery compared to perturbations in later lactation stages. This can partly be explained by a higher vulnerability to clinical health events in the beginning of lactation, while long-lasting subclinical health events might manifest clearer in later lactation stages (Burvenich et al., 2003; Leblanc, 2020). Another explanation for slowly developing perturbations in later lactation stages is the increased occurrence of contagious intramammary infections such as S. aureus infections throughout lactation as shown by e.g. Schukken et al., (2014).

### Application Potential and Future Work

Information about the perturbation characteristics can be used for several purposes, including precision phenotyping of individual cows for e.g. resilience and robustness (Elgersma et al., 2018; Berghof et al., 2019). This can be a first step towards new genetic evaluation methods in which not only production levels, but also specific lactation dynamics can be taken into account (Poppe et al., 2020). Moreover, the methodology to evaluate the perturbations’ shapes can be used to identify animals that recover easily, and thus, are less affected by external challenges (Gross et al., 2020). Characterization of perturbations is therefore a way of compiling available data into useful information for farmers and veterinarians based on available data. To this end, looking further into correlations between the perturbations’ characteristics caused by specific events can be interesting, and this provides an asset for monitoring and forecasting cure and the effect of treatment. Next to parity and lactation stage, also the effects of herd, farm, seasonal effects and diverse management practices on the characteristics and milk losses during perturbations have to be further investigated and was identified as future work. To this end, not only the specific dynamics within a type of disease, but also individual and herd mate baselines can be taken into account, as well as time series measures of complementary parameters such as milk electrical conductivity or constituents (King and DeVries, 2018). This way these data tools can be implemented for both individual and herd level monitoring applications.

## CONCLUSIONS

This study presents a new methodology to study perturbations in daily milk meter lactation curves of dairy cows managed on AMS herds. On average 3.4 perturbations were detected per lactation, associated with an average milk loss of 92.1 kg. The perturbations were characterized using a third order Savitsky-Golay smoother, and their features were calculated. A distinction between the development and recovery phase was made to allow studying specific dynamics. Mean values of the perturbation characteristics were compared between parities and lactation stage, and the differences were all found to be highly significant. Higher parity perturbations in early and mid-early lactation were shown to be more severe with faster development rates, higher milk losses and slower recovery. This study shows the potential of using high-frequency milk meter data to characterize and identify the milk yield dynamics during challenges, which can be used for applications in on-farm monitoring and precision phenotyping of complex traits such as resilience and robustness.

## ACKNOWLEDGEMENTS

Dr. Ines Adriaens received funding from a KU Leuven postdoctoral mandate grant No. PDM/19/132. The data from this study were collected in the format of farm software back-up files by the authors and the resulting database used is owned by the authors. Furthermore, this study is funded by VLAIO as an LA-trajectory ‘MastiMan’, grant No. HBC.2016.0774. Katherine Lumb (RAFT Solutions Ltd., Ripon, United Kingdom) contributed to the data collection of the UK farm data.

